# Nanopore formation in the cuticle of an insect olfactory sensillum

**DOI:** 10.1101/444729

**Authors:** Toshiya Ando, Sayaka Sekine, Sachi Inagaki, Kazuyo Misaki, Laurent Badel, Hiroyuki Moriya, Mustafa M. Sami, Yuki Itakura, Takahiro Chihara, Hokto Kazama, Shigenobu Yonemura, Shigeo Hayashi

**Affiliations:** RIKEN Center for Biosystems Dynamics Research, 2-2-3 Minatojima-minamimachi, Chuo-ku, Kobe, Hyogo 650-0047, Japan; RIKEN Center for Brain Science, 2-1 Hirosawa, Wako, Saitama, 351-0198, Japan; Department of Genetics, Graduate School of Pharmaceutical Sciences, The University of Tokyo, Hongo, Bunkyo-ku, Tokyo, Japan; Department of Biological Science, Graduate School of Science, Hiroshima University, Higashi-Hiroshima, Hiroshima 739-8526, Japan; Department of Biology, Kobe University Graduate School of Science, Kobe, Hyogo 657-8501, Japan

## Abstract

Nanometer-level patterned surface structures form the basis of biological functions including superhydrophobicity, structural coloration, and light absorption [1-3]. In insects, the cuticle overlying the olfactory sensilla has multiple small (50–200-nm diameter) pores [4-8], which are supposed to function as a filter that admits odorant molecules, while preventing the entry of larger airborne particles and limiting water loss. However, the cellular processes underlying the patterning of extracellular matrices into functional nano-structures remain unknown. Here we show that cuticular nanopores in *Drosophila* olfactory sensilla originate from a curved ultrathin film that is formed in the outermost envelope layer of the cuticle, and secreted from specialized protrusions in the plasma membrane of the hair forming (trichogen) cell. The envelope curvature coincides with plasma membrane undulations associated with endocytic structures. The *gore-tex/Osiris23* gene encodes an endosomal protein that is essential for envelope curvature, nanopore formation, and odor receptivity, and is expressed specifically in developing olfactory trichogen cells. The 24-member *Osiris* gene family is expressed in cuticle-secreting cells, and is found only in insect genomes. These results reveal an essential requirement for nanopores for odor reception and identify *Osiris* genes as a platform for investigating the evolution of surface nano-fabrication in insects.

## Results and Discussion

The great success of insects in spreading throughout nearly the entire terrestrial ecosystem has been largely due to their exoskeleton, which enabled both their adaptation to harsh environments and protection from predators. The adaptation of olfaction to airborne odorants was also key in helping insects search for food, mates, and other environmental cues, and to establish social communication [9]. Insect sensilla consist of one to several neurons and usually three supporting or auxiliary cells. The neuron(s) innervate a specialized cuticular apparatus, which often has the shape of a peg or hair and is secreted by the auxiliary cells. Later in development, these cells take over other functions, e.g. secretion of the sensillum lymph that surrounds the dendrites of the neurons. In olfactory sensilla the wall of the cuticular hair shows numerous pores which are absent from sensilla serving other modalities. For volatile odorants to reach the dendrites, numerous pores ranging in diameter from 50 to 200 nm are formed in regular arrays on the cuticles of the olfactory sensory organs [4–8]. Beneath the pores, filaments called pore tubules extend to the interior lymph and dendritic nerve endings[7,8]. These nano-scale structures are thought to allow the diffusion of small odorant molecules (0.5 to 5 nm in diameter) to reach the olfactory neurons, while preventing the entry of larger (100 to 1000 nm) airborne particles and minimizing the loss of inner lymph liquid. This selective filter system protects the olfactory neurons, which are not replaced throughout the adult life, from chemical and infectious damage. The molecular mechanisms determining the size and spatial patterns of nanopores, and how they are formed during cuticular scaffold assembly, are poorly understood. To elucidate the biological processes leading to this nanometer-order fabrication of extracellular matrix, here we investigated the molecular mechanisms of olfactory nanopore formation in the fruit fly *Drosophila melanogaster*.

Among the olfactory sensilla on the antennae and maxillary palps of *Drosophila* (Fig. 1A), we focused on those of the maxillary palp because of its simple structural composition: all ~60 of the olfactory sensilla deployed on a maxillary palp are morphologically categorized as middle length-type sensilla called basiconic sensilla [8,10]. To understand when and how the pores are formed, we studied the process of cuticle secretion in olfactory hair cells using transmission electron microscopy (TEM). On the surface of these sensilla, pores of about 50-nm diameter are distributed along the long axis at 150-to 170-nm intervals (Fig. 1B, E). These sensilla form during pupal development from sensory organ precursor (SOP) cells, which give rise to multiple SOP-cell subtypes through several rounds of asymmetric cell division [11]. Of these SOP-cell subtypes, a single cell called a shaft cell (trichogen cell) extends a cell protrusion filled with microtubules and actin filaments toward the outside of the epithelium (Fig. 1F, 44 h APF [after puparium formation], green arrowheads, [12–14]). Then, the olfactory neurons project dendrites into the olfactory sensillum sheath (Fig. 1F, 52 h APF). At 74 h APF, the cellular projection of the shaft cell retracts. The resulting shaft is now a chitin-rich cuticular sheath filled with dendrites in its inner space (Fig. 1G, 74 h APF). Adjacent to the olfactory sensilla, highly actin-rich hair cells (spinules, sp), which lack neurites, undergo a similar extension process (Fig. 1C, Fig. 1F, magenta arrowhead).

**Figure 1.**
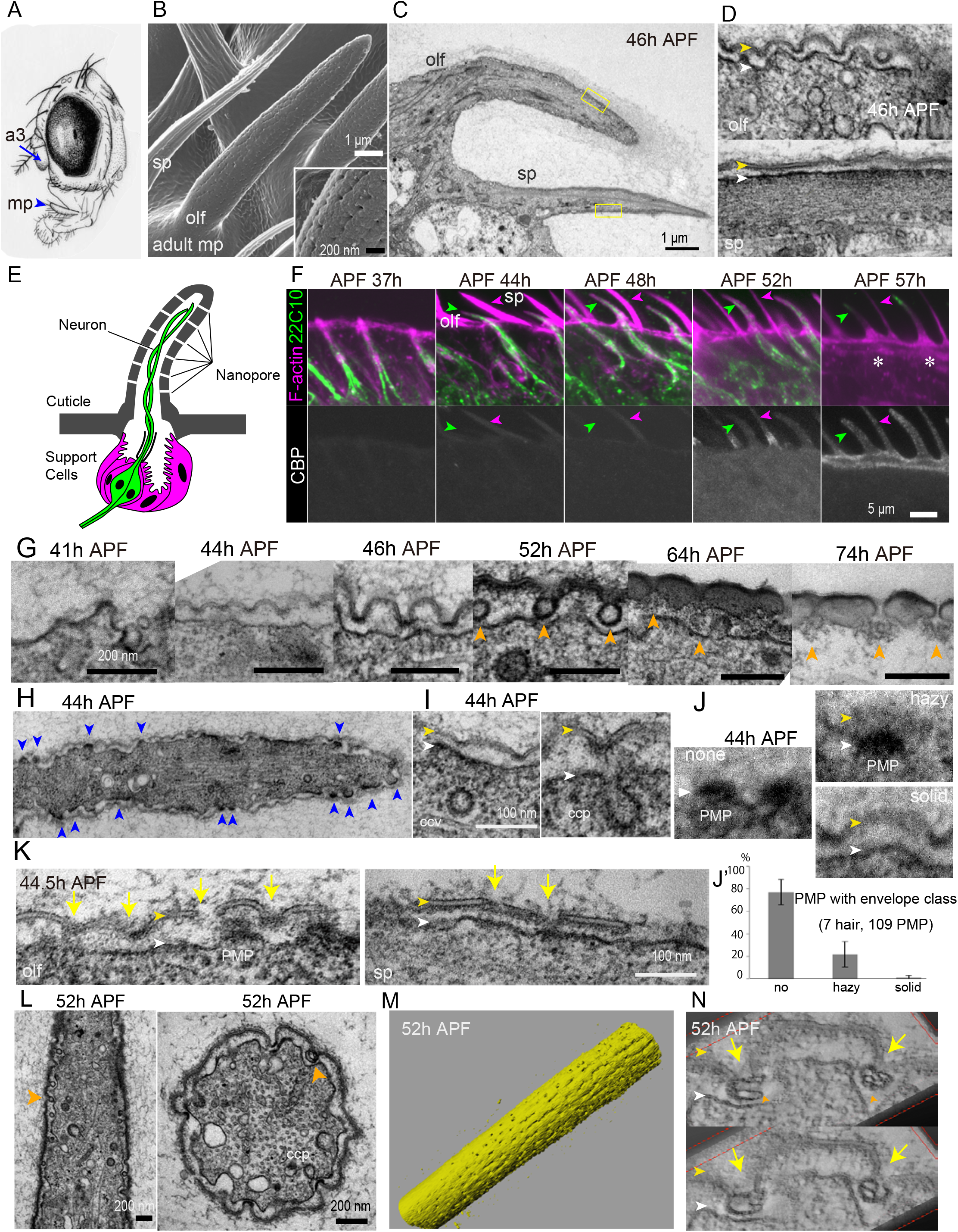
Development of olfactory nanopores. (A) Olfactory sensilla are present on the 3^rd^ antennal segment (a3) and the maxillary palp (mp) of the adult head. (B) A surface image of adult maxillary palp showing olfactory sensilla (olf) and non-sensory spinule (sp). Inset shows regularly arranged nanopores. (C) A TEM image of olf and sp at 46 h APF. (D) Enlarged crosssectional views of newly formed cuticular envelope (yellow arrowheads) and plasma membrane (white arrowheads). Note undulated envelope and plasma membrane with numerous intracellular and extracellular vesicles at the olfactory sensilla (top). In the spinule, the envelope and plasma membrane were flat, and extensive actin filaments were associated with the plasma membrane (bottom). (E) Schematic of a porous olfactory sensilla. (F) Time course of olfactory sensilla and spinule formation in maxillary palp stained for a neuronal marker (22C10, green), F-actin (magenta), and chitin (black and white panels). Neurons were associated with olfactory sensilla (green arrowhead), but not with spinule (magenta arrowhead), which was enriched in F-actin. Cuticles thickened after 52 h APF, and restricted the internal staining of neurons (*) at 57 h APF. (G) Time course of cuticle nanopore formation. Orange arrowheads show envelope-associated structures at 52 h APF, and pore tubules at 64 h APF and later. Shaft cell retraction was complete at 74 h APF. (H) Overview of olfactory sensilla at 44 h APF with extensively undulated surfaces and sporadic PMPs (plasma membrane plaque, arrowheads). (I) Examples of clathrin-coated vesicle (ccv) and clathrin-coated pit (ccp) near the site of envelope formation. (J) Three steps of envelope formation (none, hazy, solid) and the frequency of PMPs associated with each class (J’). (K) An enlarged view of olfactory sensilla and spinule cell surface associated with envelope pieces of regular size (yellow arrowheads) on top of plasma membrane (white arrowheads). Yellow arrows indicate connection points of envelope pieces. (L) Longitudinal and crosssectional views of olf at 52 h APF, showing envelope curvature and an extracellular envelope-associated structure (orange arrowhead). (M) A reconstructed surface view of olfactory sensilla obtained by SBF-SEM. (N) Complex envelope-associated structures shown by electron tomography. Top and bottom images show the same specimen viewed from different angles (See also Figure S1 and Video S1).

Insect cuticles are organized in a layered structure consisting of an envelope, epicuticle, and chitin-rich procuticle, which are secreted in a step-wise manner [15]. The first sign of envelope formation in the olfactory sensilla was observed at ~41 h APF as the appearance of electron-dense protrusions, similar to previously described plasma membrane plaques (PMPs) [16] the undule [17,18], or pimples [19] (Fig. 1G, 41 h APF). Above the PMP-positive cell surface at this stage, no distinct extracellular structure was observed (Fig. 1G, 41 h APF). Within the next few hours, each PMP was covered with diffuse material that acquired a solid appearance (Fig. 1H, J). This material eventually became the thin film of trilamellar envelope structures, and the PMP disappeared (Fig. 1D, Fig. 1G, 44, 46 h APF). PMPs appeared asynchronously over the surface of the olfactory shaft cells (Fig. 1H), and the envelope initially appeared as fragments (Fig. 1K). The piece-by-piece formation of envelope from fragments was also observed in adjacent hair cells (Fig. 1K), as previously described for the cuticles of *Drosophila* wing hair cells [17,19] and larvae [18], which form seamless cuticles. These observations imply that the PMPs function as a scaffold for the envelope-layer formation in olfactory shaft cells. Remarkably, the initial envelope pieces on the olfactory shaft cells were convex in shape, in contrast to the straight appearance of the spinule cells (Fig. 1D, arrowhead). The convex envelope pieces were connected together with diffuse materials at their lowest point (Fig. 1K, yellow arrows). The cuticles had a wavy appearance in both longitudinal and transversal sections of the olfactory shaft cells at 52 h APF (Fig. 1L), and 3D reconstruction of the nascent envelope layer using serial block face-scanning electron microscopy (SBF-SEM) revealed numerous indentations with similar sizes and patterns as the pores on the surface of the adult olfactory sensillum (Fig. 1M), suggesting that the curved ultrathin envelope layer forms the nanopores at 52 h APF.

Olfactory hair cells at 44 h APF are rich in various membrane structures, including clathrin-coated vesicles (ccv) of about 50-nm diameter and clathrin-coated pits (ccp, Fig. 1I) that appear near the plasma membrane. Image analyses revealed that 84% of the ccv (N=31) and 77% of the ccp (N=13) were located between neighboring PMPs or envelope fragments. In contrast, in the spinule cells at 44 to 46 h AFP, vesicle structures were rarely observed near the plasma membrane, where dense arrays of actin filaments occupied the space (Fig. 1D, K).

At 52 h APF, electron-dense extracellular structures appeared at the lowest point of the wavy envelope layer (Fig. 1G, 52 h APF, orange arrowheads). These structures were aligned at 150- to 200-nm intervals in sagittal sections (Fig. 1L, left) and in horizontal sections (Fig. 1L, right). Three-dimensional imaging by electron tomography revealed complex folded structures associated with the envelope layer (Fig. 1N, movie S1). These structures, which were detergent sensitive (Fig. S1), disappeared after 64 h APF and were replaced by cloudy material beneath the nascent pore structures that persisted after the shaft cell retraction (Fig. 1G, 74 h APF, orange arrowheads). It is possible that the lamellar structures are the material for the pore tubules that are thought to serve as a channel for the delivery of odorant molecules to the dendrites of the olfactory neurons [7,20].

These TEM studies suggested that a specialized cellular process for apical extracellular matrix assembly must be present in the olfactory shaft cells to produce the curved envelope pieces and nanopores. To identify the molecular components specifying the nanopores, we made the assumption that the putative nanopore-formation genes must fulfil the following criteria: 1) expressed in olfactory shaft cells at the time of shaft cell envelope formation (2 days APF), 2) but not expressed in the pore-less mechanosensory hair cells or neuronal cells, and 3) encode membrane-associated proteins. We then performed RNA-seq analysis focusing on three genetic conditions: wild type, transformation of shaft cells to neurons [overexpression of *numb*, [21]] and conversion of olfactory sensory organs into mechanosensory organs *[amos* mutants, [22]]. Pupal antennae at 52 h APF were collected for analysis (Materials and Methods, Data S1). The RNA-seq profiles of the above two genetic conditions were compared with that of wild type in triplicate samplings, and transcripts that were downregulated in the mutants were further filtered for the presence of a signal peptide or transmembrane domains. Twenty-six candidate genes selected from this screen were then knocked down in developing olfactory shaft cells by a transgenic RNAi method, and the olfactory sensillum morphology in the maxillary palp was observed by field emission scanning electron microscopy (FE-SEM). Among the 30 RNAi strains tested, two caused a loss of the nanopore phenotype (Fig. 2A, B). These two RNAi strains were targeted to the same gene (CG15538), which was named *gore-tex (gox).* According to the developmental RNA-seq profile in the modENCODE database, the *gox* transcript is detected only in 2-day-old pupae. Two *gox* mutations created by genome editing (Fig. S2) were homozygous viable and fertile, with no visible defect in adult external morphology at the macroscopic level. However, high-resolution helium ion microscopy (HIM) of the olfactory sensilla revealed a significant loss of nanopores, and this phenotype was partially rescued by two copies of HA-tagged *gox* genomic construct (gHAgox, Fig. 2A, B). Recording of the local field potential in maxillary palps revealed that the loss of *gox* by mutations or RNAi greatly reduced the responses to both strong (a mixture of odors that strongly activate sensory neurons in the palp) and weak (milder solvent) odor stimuli (Fig. 2C, D). TEM analysis of 52 h APF *gox*^1^ mutant olfactory sensilla exhibited a flat cuticular envelope and no pore structures, suggesting that the *gox* function is required to introduce curvature into the newly formed envelope fragments, prior to nanopore formation (Fig. 2E).

**Figure 2.**
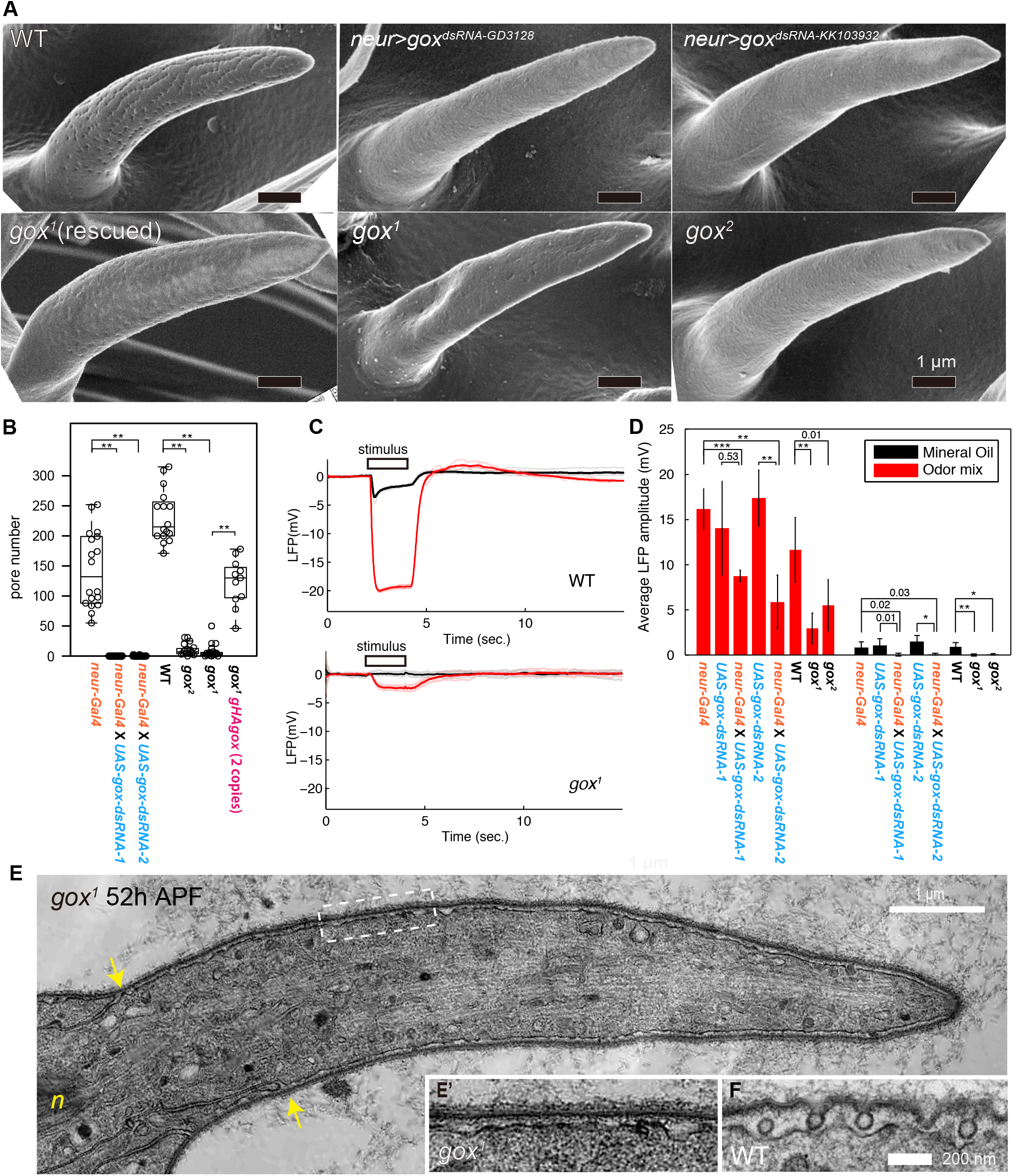
Characterization of *gore-tex* function. (A) Surface views of adult olfactory sensilla. (B) Pore counts per sensillum. Statistical significance tested by Mann–Whitney U test with Bonferroni correction is indicated by ** (p < 0.01/5). (C) Recording of local field potential (LFP) in a maxillary palp upon the application of an odor mixture (red line) and solvent only (mineral oil, black line). Solid lines are an average of 5 trials. (D) Quantification of the odor response measured from 6 recordings for each genotype. Statistical significance tested by Student’s t-test with Bonferroni correction is indicated by * (p < 0.05/12), ** (p < 0.01/12) and *** (p < 0.001/12). Concrete p-values are indicated when they are statistically nonsignificant. (E) A TEM image of a *gox*^1^ olf at 52 h APF. Arrows indicate the boundaries between the trichogen (hair) and the adjacent tormogen (socket forming) cell. n: neurite. (E’) Enlarged view of envelope in E. (F) Corresponding region of a control olf.

The predicted Gox protein contains an N-terminal signal peptide, two copies of a putative transmembrane domain, and a domain with similarity to a sequence called “domain of unknown function” (DUF1676) found in insect genomes (Fig. 3A). Gox protein expressed in *Drosophila* S2 cells was co-localized with HRS (early endosome marker) and Rab7 (late endosome marker), but not with an endoplasmic reticulum marker Calnexin (Fig. S3B). gHAgox showed Gox expression in the shaft cells of olfactory sensillum, but not in spinules (Fig. 3B, B’), and not in neurons (Fig. 3B’’). Super-resolution imaging revealed that HA-Gox was localized to intracellular vesicles that overlapped with HRS and Rab7 (Fig. 3C, D), and a subset of HA-gox was found closely associated with plasma membrane (Fig. 3E). These results indicated that Gox is a transmembrane protein that mainly localizes to the endosomes of shaft cells of olfactory sensillum.

**Figure 3.**
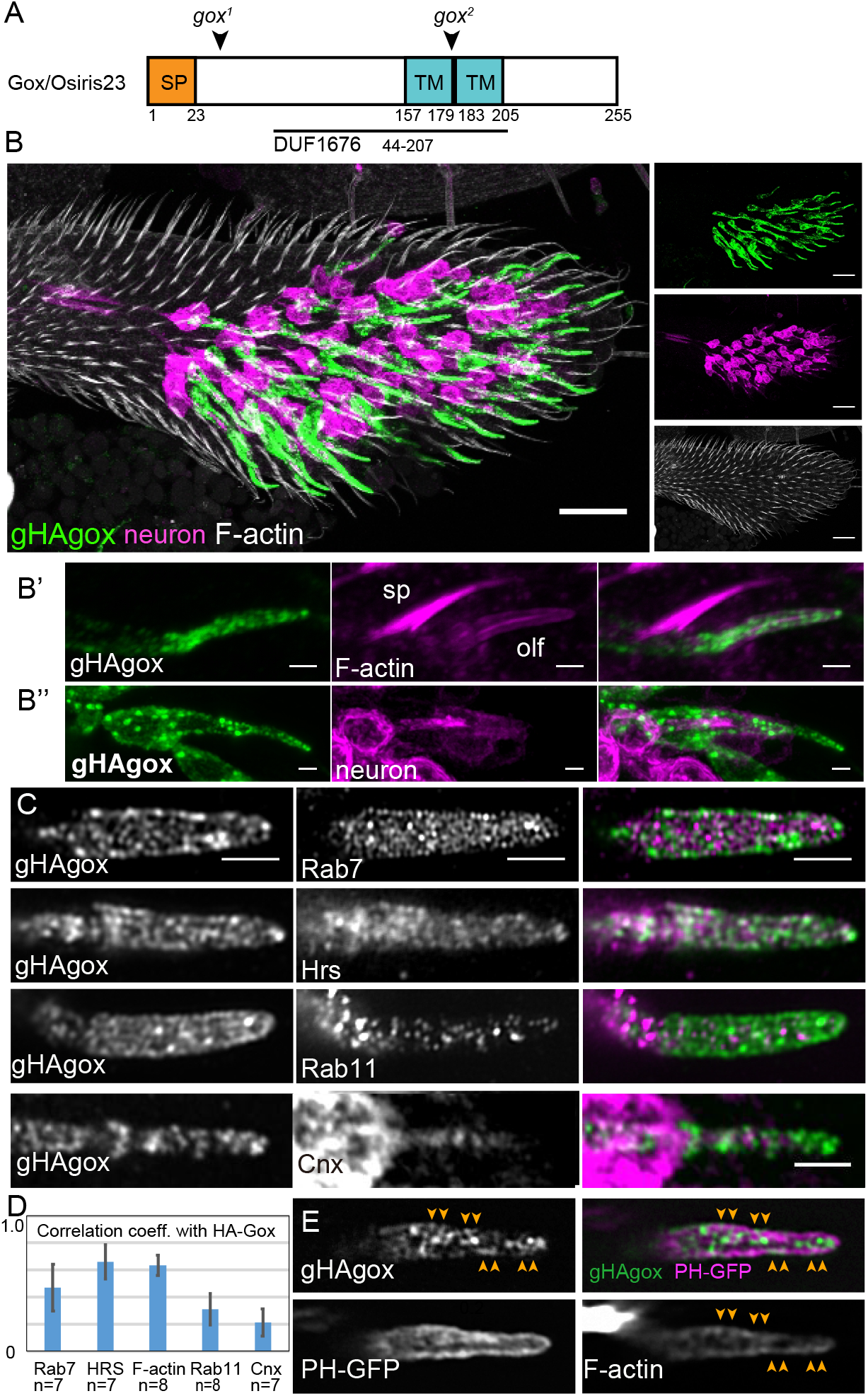
Expression of Gore-tex protein. (A) A schematic diagram of the Gore-tex domain organization (see also Figure S2). Tandem transmembrane helices predicted by multiple programs are indicated[34] (B) Whole-mount staining of a maxillary palp at 44 h APF showing the expression of an HA-tagged genomic *gox* transgene (green), neurons (magenta), and F-actin staining (gray). (B’, B’’) gHA-gox is expressed in shaft cell of olfactory sensillum (olf) with a low level of F-actin, and not in spinule (sp) or neurons that innervate into olfactory sensillum. (C) Subcellular localization of gHA-Gox and organelle markers (Rab7, Hrs, Rab11, Cnx, see Fig. S3 for expression in S2 cells). (D) Colocalization analysis of HA-Gox and organelle markers. (E) Colocalization of HA-gox, plasma membrane (PH-GFP) and F-actin (arrowhead) markers. Scale bar: 20μm (B), 2μm (B’, B’’, C).

The *gox* gene belongs to a 24-member gene family called *Osiris (Osi).* Of the 24 *Osiris* genes, 21 are clustered at position 83E of chromosome 3R, and are included in a locus that exhibits a rare dosage sensitivity (triplo-lethal and haplo-lethal) [23,24]. The other three is located at 32E *(Osi21)*, 87E *(Osi22)*, and 99F *(gox/Osi23).* To obtain information about the function of *Osiris* genes, we investigated the mRNA expression of 24 *Osiris* genes in *Drosophila* embryos by *in situ* hybridization. We found that all of the *Osi* genes were expressed at variable levels and patterns in late embryonic stages when the cuticle is secreted, in various tissues, including the epidermis, mouth parts, foregut, hindgut, trachea, and spiracles (Fig. 4A, Table S1, Fig. S4, supplementary text, see also [25,26] for a subset of *Osi* gene expression in trachea). We classified the *Osiris* expression patterns into 8 types, which collectively covered essentially all of the cuticle-producing organs of first-instar larvae (Fig. S4). Of these mRNAs, *Osi6* and *Osi7* were expressed broadly in the epidermis and head skeleton. When *Osi6* and *Osi7* were knocked out by genome editing, the resulting heterozygous animals were weak and sluggish, and homozygous animals showed poor formation of the cepharo-pharyngeal skeleton and denticle belts, resulting in complete *(Osi6)* or partial *(Osi7)* failure of hatching (Fig. 4B, C, Supplemental video S2 and S3). Homozygous *Osi19* mutants were semi-lethal and showed a loss of ocelli and a rough-eye phenotype (Fig. 4E). These results collectively indicated that *Osiris* genes are expressed mainly in cuticle-secreting epidermal tissues at cuticle-secretion stages, and that the available null mutations of each four of them (*Osi6, Osi7, Osi19*, and *gox/Osi23)* caused unique defects in cuticle patterning or functions, as previously suggested from bioinformatics analysis [27].

**Figure 4.**
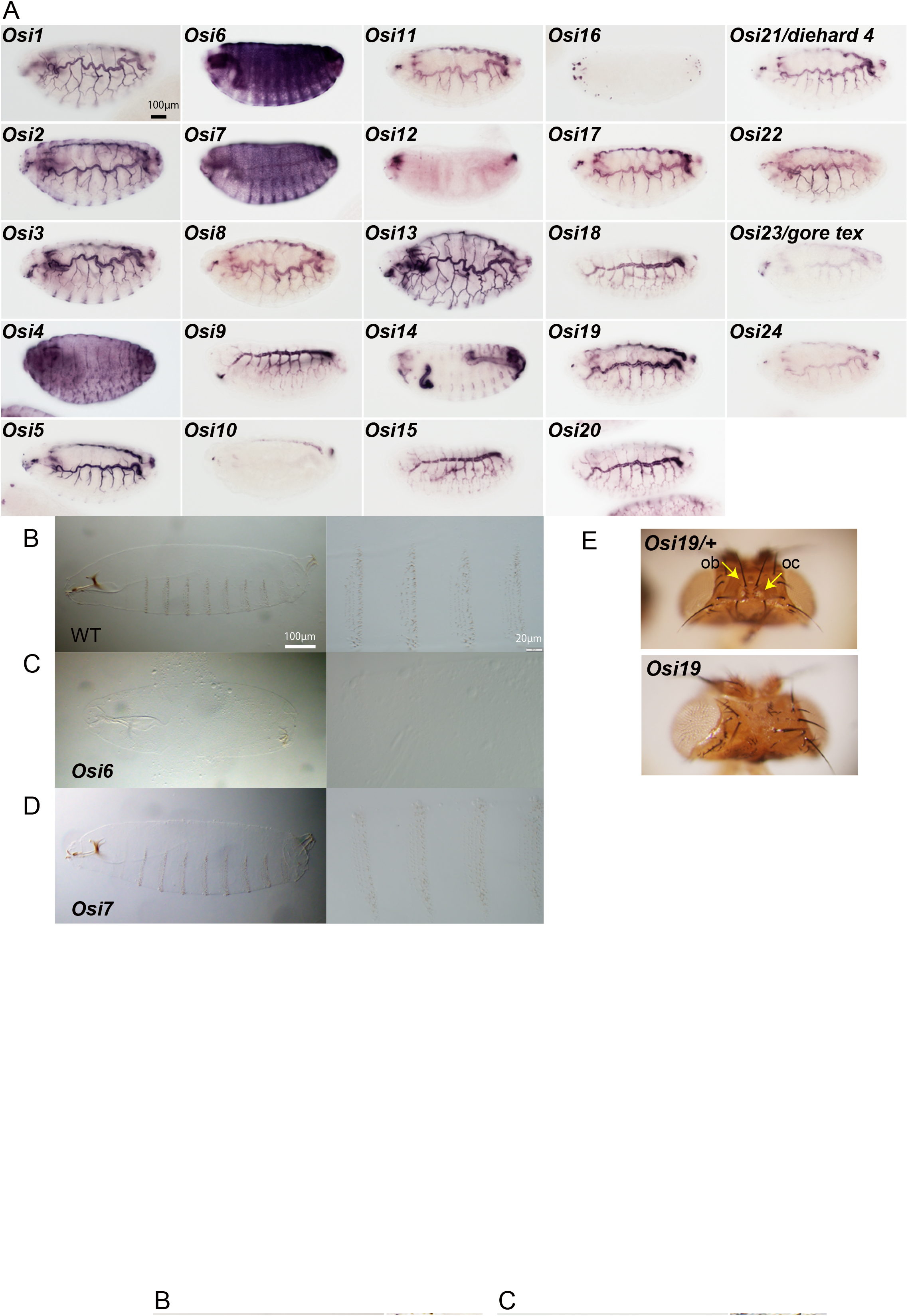
Expression and function of the *Osiris* gene family. (A) Embryonic mRNA expression patterns of 24 *Osiris* genes. See Table S1 and Fig. S4 for more detailed description. (B-D) Cuticle preparation of control Oregon R (B), *Osi6* (C) and *Osi7* (D) embryos (Video S2 and S3 show their hatching behavior). Images of entire embryo (left) and denticle belts (right). *Osi6* embryos have very weak and diminutive cuticles. (E) Adult heads of *Osi19* mutants. Ocelli (oc) were missing, and orbital bristles (ob) were partially lost in homozygotes.

*Osiris* gene families with DUF1676 domain have been found in a number of insect genomes, but not in the genome of other animal classes including Crustaceans, Myriapod, Chelicerata and Entognatha (Collembola) [27,28], suggesting that this gene family was acquired at an early stage of insect genome evolution, when crustacean-like insect ancestors changed their habitat from an aquatic to a terrestrial environment and expanded their niche [29]. This finding leads to an interesting hypothesis that the acquisition of *gore-tex/Osiris23* drove the change in olfactory sensillum structure from the thin, permeable cuticles of crustaceans to the porous structures of insects [30]. How the Gore-tex protein, which is localized to endosomal compartments, affects the extracellular events of cuticle envelope assembly is still unclear. One possibility is that Goretex regulates the trafficking of specific effector molecules for envelope assembly such as Trynity[31], as previously suggested for the function of *Osi21/die4* in regulating endocytosed rhodopsin in photoreceptor cells [32]. Alternatively, Gore-tex protein may help to pattern the endocytosis of the olfactory shaft cell membrane. It is notable that the spacing and alignment pattern of the nanopores is highly regular, coinciding to the position of clathrin coated pit and clathrin coated vesicle adjacent to the PMPs at ~44 h APF. Targeting the site of endocytosis between PMPs would enhance the periodicity of endocytic and exocytic activities, leading to cuticle undulation, as previously suggested for the taenidial fold formation in the *Drosophila* trachea [33]. The nano-scale surface patterning of extracellular matrices forms the basis for the vast variety of functional biological surfaces with specific properties of, for example, structural color, water repellency, and light absorption [1–3]. Further studies of *gore-tex/Osiris23* and its family members should uncover mechanisms behind the specialized features of biological nano-patterning in the apical extracellular matrix.

## Acknowledgments

We thank Shigehiro Kuraku and his laboratory members for technical support for the RNA-seq analysis; Carl Zeiss Japan, and Shinichi Ogawa (the National Institute of Advanced Industrial Science and Technology) for the use of HIM; Atsushi Yamaguchi (Hyogo Prefectural Institute of Technology) for the use of FE-SEM; Andrew Jarman (The University of Edinburgh), Masayuki Miura (The University of Tokyo), the *Drosophila* stock centers of Kyoto, NIG, and Bloomington for fly stocks; the Developmental Studies Hybridoma Bank, Akira Nakamura (Kumamoto University), and Hugo Bellen (Baylor College of Medicine) for antibodies; Hosei Wada for genome engineering support; Hiromi Niwa for illustration; and Mai Shibata for support for cell culture experiments. We are grateful to Masayuki Miura (The University of Tokyo) for supporting the project performed by H.M. and T.C., and Rudolf Alexander Steinbrecht (Max Planck Institute for Ornithology) for discussion. SS is a JSPS Research Fellow. A part of this work (HIM) was conducted at the Nano-Processing Facility, supported by IBEC Innovation Platform, AIST. This study was supported by JSPS KAKENHI Grant Number 15K18811 to TA and, 18J40165 and 18K14746 to SS.

## Author contributions

T.A. and S.H. conceived of the project, and designed the experiments. T.A. performed the TEM analysis with K.M. and S.Y., and the electrophysiological analysis with L.B. and H.K. S.S. performed the cellular localization analysis of *gox/Osi23* with the help of S.H., and created the *gox/Osi23* mutants. The other *Osi* mutants were created by H.M. and T.C. and were analyzed by S.I. and Y.I. S.I. performed the *in situ* hybridization analysis. M.M.S. developed the tool for the image analysis. T.A. performed the other experiments, and analyzed the experimental data with the help of S.H. T.A and S.H. wrote the manuscript with the help of S.S. and H.K.

## STAR Methods

### CONTACT FOR REAGENT AND RESOURCE SHARING

Further information and requests for resources and reagents should be directed to and will be fulfilled by the Lead Contact, Shigeo Hayashi.

### EXPERIMENTAL MODEL AND SUBJECT DETAILS

*Drosophila melanogaster* strains were maintained in plastic vials with standard yeast-cornmeal-agar media at 25°C. Female adult flies within 1 week after eclosion were used for analysis of external morphology by helium ion microscopy or scanning electron microscopy. For analysis of pupal tissues, white prepupae were collected (APF 0h) and left to develop at 25°C for 2-3 days until dissection. Typically, single pupa was analyzed for each time point of TEM analysis. Some variation in developmental time among population and between different genotype and sex is possible.

### METHOD DETAILS

#### Transmission electron microscopy (TEM)

The method previously described for analysis of mechanosensory bristles was adopted[35]. Head tissues were dissected from pupae in phosphate-buffered saline (PBS), and immediately transferred to fixation buffer (2.5% glutaraldehyde, 2% formaldehyde, 0.1 M cacodylate buffer, pH 7.4) on ice. Pupal cuticles were removed in the buffer. The dissected tissues were stored at 4 °C for more than one night. After fixation, the tissues were washed three times for 10 min each in 0.1 M cacodylate buffer, and post-fixed in 1% OsO_4_ in 0.1 M cacodylate buffer for 2 h on ice in the dark. The fixed tissues were then washed three times for 10 min each in water, and stained en bloc in 0.5% uranyl acetate aqueous solution overnight at room temperature. The tissues were subsequently dehydrated in a graded ethanol series (50, 70, 80, 90, 95, and 99.5%) for 10 min at each concentration, and transferred to 100% ethanol for two 20-min incubations. After dehydration, the tissues were incubated in propylene oxide for 20 min twice, infiltrated with a 1:1 mixture of propylene oxide/QY1 (n-butyl-glycidyl-ether), incubated in 100% QY1 for 20 min, infiltrated with a 1:1 mixture of QY1/resin (Quetol 651, NEM) for 2 h, and finally placed into 100% resin. The resin was polymerized for more than 48 h at 60 °C. After trimming the embedded tissues other than the anterior head region including the mouthparts and maxillary palps, semi-thin sections (400–700 nm) were cut from the anterior side of the head. After images of maxillary palps started to appear in the sections, ultrathin sections (approximately 50 nm) were cut and mounted on 50-mesh formvar-coated copper grids. Sagittal sections of olfactory shaft cells were obtained by cutting parallel to the dorsal plane of a maxillary palp. Coronal sections of olfactory shaft cells were obtained by cutting perpendicularly to the proximodistal axis of a maxillary palp. The sections were stained with 4% aqueous uranyl acetate in the dark for 10 min, and with Reynold’s lead citrate for 3 min. Observations were performed using a JEM-1010 transmission electron microscope (JEOL) at 100 kV accelerating voltage, and images were captured by a Gatan Bioscan 792 digital camera. Following number of animals and sensillum were analyzed for wild type morphology (Oregon R) in each stage: 41h APF (1 animal, 5 sensillum), 43.5 APF (1, 7), 44.5h APF (1, 7), 46h APF (1, 7), 52h APF (3, 16), 64h APF (1, 2), 70h APF (1, 1), 78h APF (1, 1).

#### Electron tomography (ET)

Sixty-nanometer sections cut from blocks prepared for TEM were used for electron tomography analysis. Typically, the sample grid was sequentially tilted in the range of +/− 70° and images were taken using a JEM-1400 Plus equipped with Recorder software (JEOL). The obtained image series were 3D reconstructed and analyzed with Composer and Visualizer-Kai software (JEOL).

#### Scanning electron and helium ion microscopy (SEM & HIM)

Adult flies of each genetic background were collected into 70% ethanol, and stored at room temperature until use. Head tissues were collected in PBS, then fixed and dehydrated as described for the TEM analysis. After dehydration, the samples were dried in a vacuum, mounted on double-sided carbon tape placed on a brass pedestal, and coated with OsO_4_ at approximately 5-nm thickness using an osmium coater (Neoc-Pro, Meiwafosis). Observations were performed using a field emission scanning electron microscope (JSM-7610, JEOL) or a helium ion microscope (ORION NanoFab, Carl Zeiss).

#### Serial block face scanning electron microscopy (SBF-SEM)

Pupal heads at 52 h APF were prefixed as described for TEM, and postfixed with 2% OsO_4_, 1.5% (wt/vol) potassium ferricyanide, 2 mM CaCl_2_, and 0.15 M sodium cacodylate buffer (pH 7.4). After being washed with cacodylate buffer, the samples were placed in 3% low-melting agarose, then dehydrated and embedded as described for TEM. Serial block face images were taken with a MERLIN VP FE-SEM (Carl Zeiss) equipped with a 3View^®^ ultra-microtome (Gatan, Inc.). The obtained images were aligned using a custom ImageJ plugin (CoordinateShiftHyper) (Hosei Wada), and image registration was performed using the StackReg imageJ plugin [36]. After contrast of the images was enhanced to extract the surface structure, 3D volume data of a developing olfactory sensillum was reconstructed using the Imaris software (Bitplain).

#### Fluorescence microscopy

Super-resolution images of the olfactory sensilla and spinules on the maxillary palp were acquired on an inverted Zeiss LSM 880 confocal microscope with an Airyscan detector, and a Plan-Apochromat 63x/1.4NA oil immersion objective lens or a Plan-Apochromat 40x/1.3 oil immersion objective lens (for whole maxillary palp). The images were reconstructed by Airyscan processing with the 3D or 2D auto setting of the Zen software (Carl Zeiss). Imaging data were analyzed with imageJ/Fiji.

For sample preparation, after selecting the appropriate pupal stage, the head was exposed from the pupal case and immediately pre-fixed with 4% paraformaldehyde, 0.3% TritonX-100 in PBS for 20 min at room temperature, then the pupal cuticle was removed and the head was fixed again with the same fixation buffer for 20 min at room temperature. The staining was performed as described previously [35], and after adding the mounting solution, SlowFade Diamond Antifade Mountant (ThermoFisher, S36972), the maxillary palp was detached from the head and mounted on a slide glass. The following antibodies and staining reagents were used: rabbit anti-Rab7 (1:100, gift from Akira Nakamura), rabbit anti-Rab11 (1:50, gift from Akira Nakamura), mouse anti-Hrs (1:10; DSHB, Hrs 27-4), rat anti-HA (1:500; Merck, 3F10), mouse anti-Calnexin99A (1:10; DSHB, Cnx99A 6-2-1) and Phalloidin-TRITC (1:100; Merck, P1951), Alexa488-labeled chitin binding probe (1:50 New England Biolab unpublished, [37]). The following secondary goat antibodies were used at 1:200: anti-rabbit Alexa 488 highly cross adsorbed (ThermoFisher; A11034), anti-mouse Alexa 488 highly cross-adsorbed (ThermoFisher; A11029), and anti-rat Alexa 647 (Jackson; 112-606-143). To quantify the colocalization between gHAgox and the organelle markers, a single section was selected and approximately three fourth of the distal region of the olfactory sensillum was cropped manually to collect the signals only from shaft cell. Then a Pearson’s R value with no threshold was calculated using Fiji’s function, Coloc2.

#### Differential gene expression analysis

We performed pupal antennal RNA-seq analysis using a wild-type strain (Canton-S), and two mutant strains (*amos*^1^ and *neur>nb*). Each strain was reared at 25 °C. In the morning from Monday to Wednesday, male white pupae were collected from rearing vials, and incubated for 52 h. In the afternoon from Wednesday to Friday, pupae at 52 h APF were collected into ice cold water, and dissected within an hour.

Dissection was performed using fine forceps in droplets of PBS on a sheet of plastic film. First, the anterior head tissues including the antennae were dissected in the first droplet. Then, the pair of antennae were dissected in the second droplet. Antennal a3 segments and aristae attached to them, were dissected in the third droplet. The collected tissues were washed in the fourth to sixth droplets. Every transfer of the dissected tissues to the next droplet was performed using the PBS micro-droplet formed between the edges of forceps. After every tissue transfer, the forceps were washed in boiling water for 10 seconds, washed in ice cold water, and wiped with a fresh paper towel. The washed tissues were collected into Buffer RLE (RNeasy Micro Kit, Qiagen) supplemented with 2-mercaptoethanol on ice, and manually crushed with a BioMasher II (Nippi). At the end of each day, the collected lysate was frozen, and stored at −80°C until use. To collect 100 ng of total RNA for each sample and three biological replicates for each strain, this procedure was repeated for a month and a half in total. In the Canton-S strain (gift from Masayuki Miura) and the *neur>nb* mutant *(y^*^, w^*^, P{ey-FLP.N}2, P{GMR-lacZ.C(38.1)}TPN1/Y; UAS-nb/+; P{GawB}neur^GAL4-A101^*, *P(αTub84B(FRT.GAL80)}3/+)*, antennae of approximately 40 flies were pooled as a single sample. In the *amos* mutants (*amos^1^, pr^1^/ Df[2L]M36F-S6*) (gift from Andrew Jarman), antennae of approximately 75 flies were pooled as a single sample.

Total RNA was extracted using an RNeasy Micro Kit (Qiagen) according to the manufacturer’s instructions. Libraries for RNA-seq were prepared using 91 ng of the extracted total RNA following the standard protocol of the TruSeq Stranded mRNA Sample Prep Kit (Illumina). They were amplified with 9 cycles of polymerase chain reaction, followed by onboard cluster generation using a HiSeq SR Rapid Cluster Kit v2 (Illumina), and sequenced on the Rapid Run Mode of Illumina HiSeq 1500 (Illumina) to obtain single-end 80-nt single reads.

The quality of the obtained reads was evaluated using the FastQC quality check package (version 0.11.3), and were processed by Cutadapt (version 1.8) and Fastx (version 0.0.14) using the default parameters to remove the Illumina TruSeq adaptor sequence and low quality reads. Ribosomal RNA sequences were filtered out by mapping the RNA-seq reads to a *Drosophila melanogaster* reference genome sequences (BDGP release 6) using Bowtie (version 2.2.5) software and the associated gene annotation file (D. *melanogaster* Ensembl release 83 annotation). Having ensured the high quality of the data, sequence reads for each library were mapped independently to the *Drosophila* reference genome sequences (BDGP release 6), using the spliced aligner Tophat (version 2.0.14) with the default parameter settings. This process yielded a high percentage of uniquely mapped reads (>92%) for all libraries.

The mRNA quantifications of each annotated gene (D. *melanogaster* Ensembl release 83) were performed using the Cuffdiff program in the Cufflinks package (version 2.2.1). To investigate the differential gene expression between fly strains, the raw RNA-seq read count data were normalized using the trimmed mean of M-values (TMM) method, and the differential expression between strains was quantified using the R package edgeR (version 3.8.6).

#### Computational prediction of transmembrane and signal peptide domains

A *Drosophila* polypeptide dataset (30,452 sequences) was downloaded from Flybase [38], and used for protein domain prediction. Signal peptide and transmembrane domains were predicted using the SignalP (version 4.1e) and TMHMM (version 2.0c) programs, respectively. The annotation of each protein was combined with the RNA-seq data. Incongruence between the Flybase IDs (FBgn ID) in March 2016 and those in the *D. melanogaster* Ensembl release 83 annotation, were semi-manually corrected using custom Perl and AppleScript scripts by accessing the Flybase webpage.

#### Molecular cloning

To clone the open reading frame (ORF) sequence of gox, template cDNA for reverse transcription-polymerase chain reaction (RT-PCR) was synthesized from the total RNA extracted from pupal antennae. The dissection of pupal antennae and total RNA extraction were performed as described for the RNA-seq analysis. Approximately 20 ng of total RNA was treated with 2 U of DNase I (TaKaRa) for 15 min at 37 °C. After heat inactivation, first-strand cDNA was synthesised using the Super Script III First-Strand Synthesis System for RT-PCR (Invitrogen). The ORF of *gox* was predicted using public RNA-seq data (modEncode), and was amplified by RT-PCR using undiluted 1st strand cDNA as a template, KOD Plus Neo (Toyobo) as the DNA polymerase, and gene-specific primers (Table S1). The amplified PCR product was cloned into the pCR4-TOPO DNA vector using the TOPO TA Cloning Kit for sequencing (Thermo Fisher Scientific), after adding an adenine residue at the 3′ end using Ex-Taq DNA polymerase (TaKaRa). The sequence of the ORF was checked by a DNA sequencing service at RIKEN CLST.

The DNA vectors for overexpression were constructed by amplifying each cloned *gox* ORF by PCR and inserting it into the pUAST-attB vector double digested with *Bgl* II and *Xho* I. All ligation reactions were mediated by the NEBuilder HiFi DNA assembly master mix. pUAST-attB-HAgox and pUAST-attB-gox.Flag carry a hemagglutinin (HA) tag and a 3XFLAG tag at the BglII site immediately downstream of the signal peptide, and the C-terminus of the *gox* ORF, respectively. The primers used are listed in Table S1.

To construct the HA-tagged genomic rescue construct, the *gox* genomic sequence including the *gox* gene and its flanking intergenic sequences (2-kb upstream and 2-kb downstream, 5 kb in total including the *gox* gene) was amplified by PCR using KOD -plus-Neo (Toyobo), with primers SS3 and SS2, using genomic DNA (Canton-S strain) as a template, and cloned into the pattB vector. The pattB/genomic *gox* vector was then cut at a Bgl*II* (Nippon Gene) site that resides right after the signal sequence, and an HA-tag containing a DNA fragment generated by an annealing two oligos (SS10 and SS11) was inserted using the In-Fusion HD Cloning Kit (Clontech). The sequence around the *gox* gene was checked by DNA sequencing.

To construct the HA-tagged genomic rescue construct, the *gox* genomic sequence including the *gox* gene and its flanking intergenic sequences (2-kb upstream and 2-kb downstream, 5 kb in total including the *gox* gene) was amplified by PCR using KOD -plus-Neo (Toyobo), with primers SS3 and SS2, using genomic DNA (Canton-S strain) as a template, and cloned into the pattB vector. The pattB/genomic *gox* vector was then cut at a Bgl*II* (Nippon Gene) site that resides right after the signal sequence, and an HA-tag containing a DNA fragment generated by an annealing two oligos (SS10 and SS11) was inserted using the In-Fusion HD Cloning Kit (Clontech). The sequence around the *gox* gene was checked by DNA sequencing.

#### Germ-line transformation and genome editing

The U6.2 promoter-driven short guide RNA (sgRNA) expression vectors used for the CRISPR/Cas9-mediated mutagenesis of *gox/Osi23* were constructed by cloning annealed oligo DNAs (gore-tex-CRISPR1 for *gox^2^* and gore-tex-CRISPR2-rzym for *gox^1^*) into the pBFV-U6.2 vector digested with BbsI [39]. Note that the target sequence choice for sgRNA expressed with the U6.2 RNA Polymerase III promoter should start with a guanine base. To overcome this limitation, we inserted a hammerhead ribozyme sequence between the transcription initiation site and the target sequence (gore-tex-CRISPR2-rzym) [40]. The guide RNA vector plasmids were injected into the *y^2^ cho^2^ v^1^ P{nos-Cas9, y^+^, v^+^}1A/FM7c, Kr-GAL4, UAS-GFP* strain [39]. The mutant flies were screened using the heteroduplex mobility shift assay [41], and the genome sequences of the homozygous mutant flies were checked by DNA sequencing.

*Osi6, Osi7*, and *Osi19* were knocked out by inserting a 3xP3 DsRed cassette into each gene according to the method of homology-directed repair [42] using the oligonucleotides listed in Table S2. Guide RNA expression vectors were constructed by inserting an annealed oligonucleotide designed against each gene into U6b-sgRNA-short[43]. Donor vectors were constructed by inserting 5’ and 3’ homology arm fragments flanking the guide RNA cleavage site of each gene into pHD-DsRed-attP using the In-Fusion PCR cloning kit (Takara). Mixtures of guide RNA and donor vector plasmids were injected into the *y^1^ M{vas-Cas9.RFP-}ZH-2A w[1118] / FM7a, P{Tb1}FM7A or w^1118^ PBac{vas-Cas9} VK00037/ CyO P{Tb1}CprCyO-A* strains by BestGene Inc. After crossing injected flies and *w^1118^*, progenies with DsRed-positive eyes were selected and verified by PCR.

#### RNA in situ hybridization

Digoxygenin-labeled RNA probes were synthesized with T7 RNA polymerase from templates that were PCR-amplified with gene-specific primers (Data S2) and purified by gel filtration. Whole-mount RNA in situ hybridization was performed according the manual procedure described previously [44,45]. Embryos were mounted in 80% glycerol and photographed using an Olympus BX53 upright microscope equipped with DIC optics and a DP74 digital camera.

#### Cell culture and transfection

*Drosophila* Schneider 2 cells obtained from the RIKEN Bioresource Center (RCB1153) were cultured in Schneider’s Drosophila Medium (1x) + L-Glutamine (Gibco 21720-024) supplemented with 10% fetal bovine serum and Penicillin Streptomycin (Gibco 15140-122). Cells cultured on glass coverslips were transfected with pUAST-HA-gox and pWA-Gal4 (http://flybase.org/reports/FBtp0010842) using Effectene (Qiagen 301425). Forty-eight hours later, the cells were fixed with 4% paraformaldehyde in PBS, blocked in 2% BSA, 0.1% Triton X-100, and stained with the set of antibodies listed in Key Resource Table. After staining, the cells were mounted in VECTASHIELD Mounting Medium (Vector Laboratories H-1200) and observed with an Olympus FV1000 confocal microscope.

#### Embryo cuticle preparation and live imaging of hatching behavior

For cuticle preparations, larvae were placed on a glass slide with Hoyer’s medium and lactic acid (Wako) 1: 1 and cover glass, and incubated at 60-65 °C overnight before observation with differential interference microscopy. For live imaging of hatching behavior, dechorionated stage-17 embryos with visible Malpighian tubules and slightly pigmented mouth cuticle placed on a plate with heptane glue (heptane with sticky tape), covered with water and imaged with a Digital Microscope VHX-6000 (Keyence) as described [31].

#### LFP recording from the maxillary palp

As the method of odor response assay LFP recording, not single sensilla recordings was chosen to avoid breaking cuticle. Flies were gently placed at the trimmed end of a plastic pipette tip. The proboscis was fixed to the edge of the pipette tip with a piece of dental wax, and a maxillary palp was immobilized with a hook-shaped glass capillary, respectively. A saline-filled reference electrode was inserted into the eye, and a saline-filled sharp electrode was inserted into a sensillum under a microscope (BX61WI, Olympus) equipped with a 50x air objective lens. The saline contained the following (in mM): 103 NaCl, 3 KCl, 5 N-tris(hydroxymethyl)methyl-2-aminoethane-sulfonic acid, 8 trehalose, 10 glucose, 26 NaHCO_3_, 1 NaH_2_PO_4_, 1.5 CaCl_2_, and 4 MgCl_2_ (adjusted to 270–275 mOsm). Signals recorded with an amplifier (Multiclamp 700B, Molecular Devices) were low-pass filtered at 2 kHz and digitized at 10 kHz. A mixture of odors that strongly activate sensory neurons in the palp (benzaldehyde, fenchone, ethyl butyrate, and pentyl acetate all diluted 100-fold) or a milder solvent (mineral oil) were applied with a custom-made olfactometer, the design of which was modified from [46]. Each olfactory stimulus was applied for 2 s while keeping the total amount of air flow constant throughout the experiment. The LFP amplitude was averaged over the odor application period and across 5 trials per fly. Mean ± sem across flies is plotted in Fig. 2D.

### QUANTIFICATION AND STATISTICAL ANALYSIS

To count the pore number, olfactory sensillum in the HIM images were segmented, and pores were identified using a custom-written pore-detection program, and manual correction software developed by MMS (MATLAB). Counting was performed blindly: one person (SH) counted images, and another (TA) performed the statistical analysis. Statistical analysis was performed using the wilcox.exact program in the R package, exactRankTests. Statistical analysis of LFP recording was performed using the ttest function in Matlab.

### DATA AND SOFTWARE AVAILABILITY

RNA seq data was deposited to Genbank (SRP155317).

### KEY RESOURCES TABLE

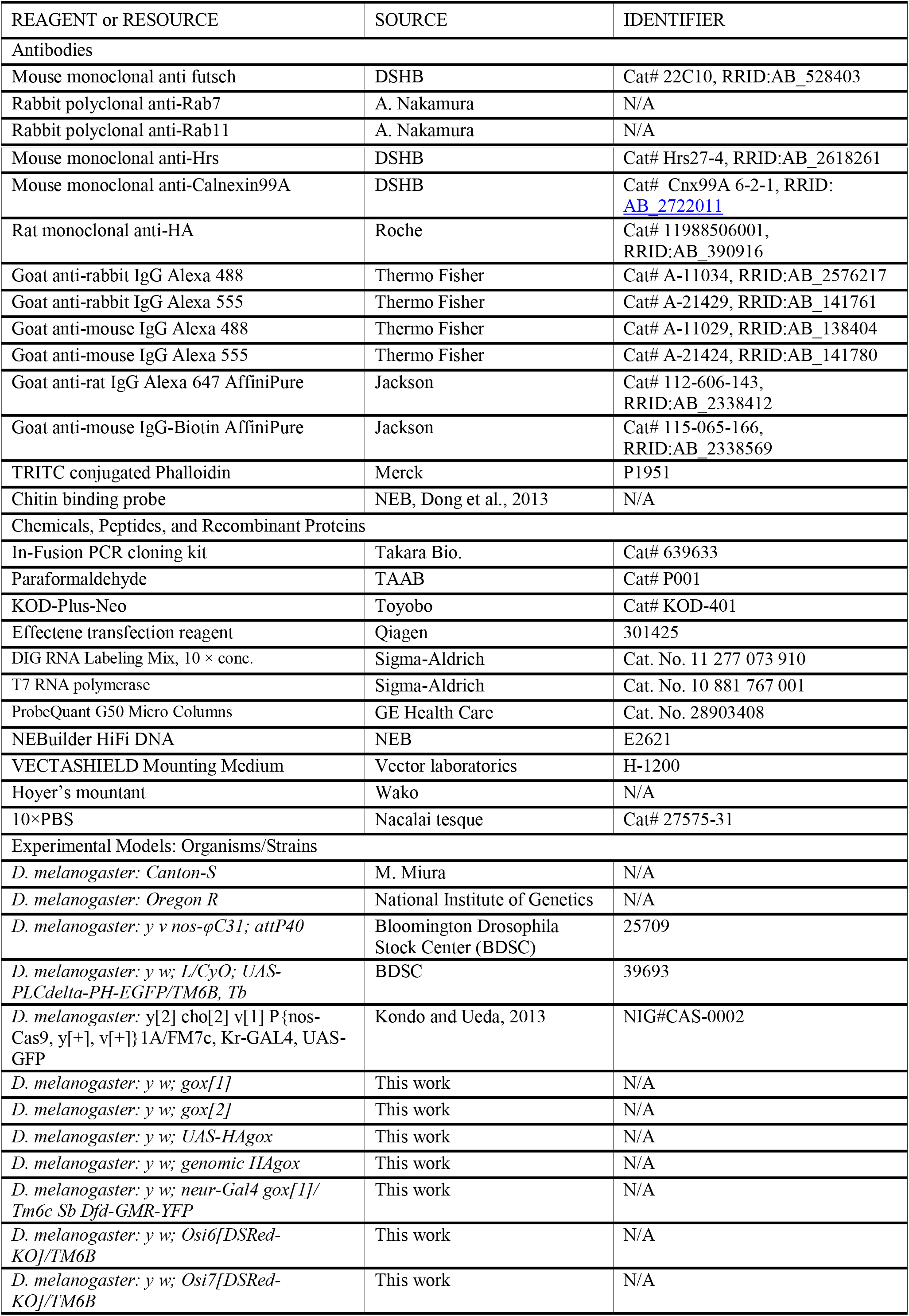

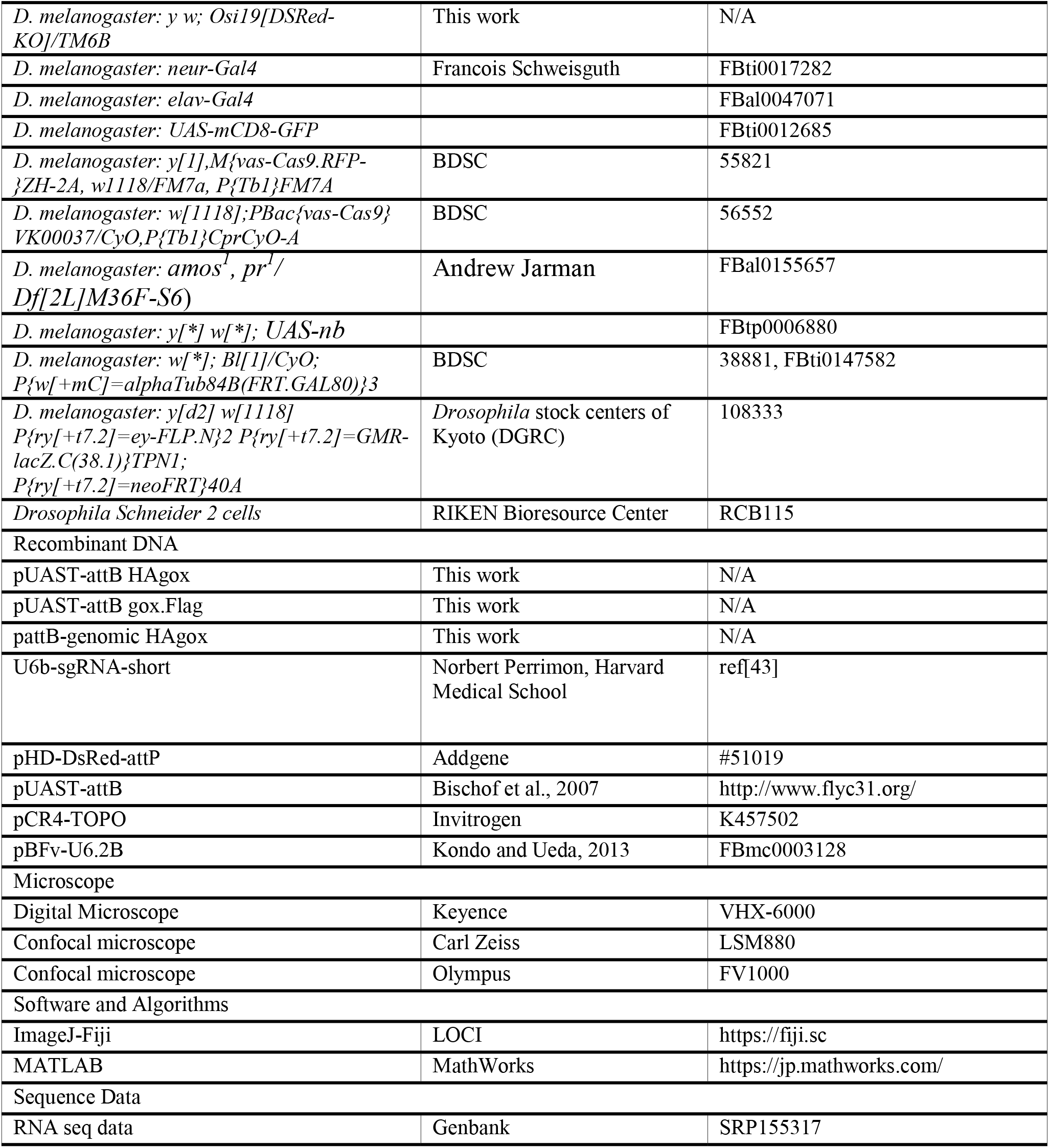

**Video S1.**

Electron tomography of the pore-forming area at 52 h APF. The sample was tilted in the +/− 70° range.

**Video S2.**

Hatching behavior of *Osi6* mutant embryos (bottom row) and control Oregon R embryos (top row) in stage 17. Control embryos hatched within 9 hours of incubation at 25°C. Although *Osi6* embryos moved vigorously with air-filled trachea, none of them hatched by 19 hours, probably due to near absence of mouse hook.

**Video S3.**

Hatching behavior of *Osi7* mutant embryos (bottom row) and control Oregon R embryos (top row) in stage 17. Control embryos hatched within 7 hours of incubation at 25°C (except for one dead embryo). Six out of 8 *Osi7* embryos failed to hatch after 12 hours.

**Data S1.**

Results of RNA-seq and bioinformatics searches and the fly strains used for the RNAi experiment.

**Data S2.**

Oligonucleotides used in this study.

