## Supplementary material for "Nanopore formation in the cuticle of an insect olfactory sensillum": Fig. S1-S4, Table S1

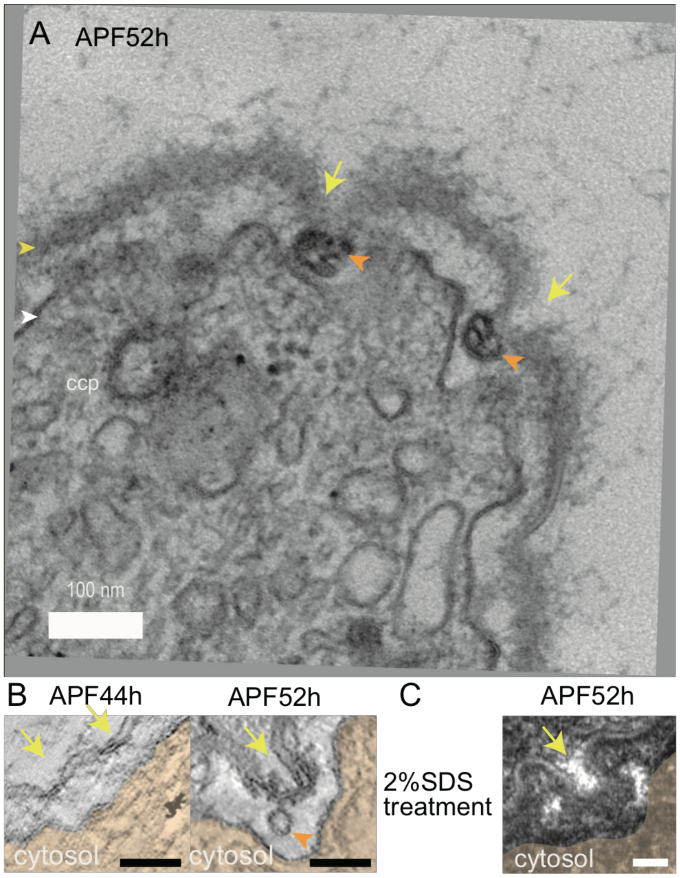


**Figure S1. Electron tomography of a developing olfactory sensillum. Related to Figure 1** (A). View of a 60-nm section corresponding to the tip of the 52 h APF olfactory sensillum used for the electron tomography analysis shown in Fig. 1M. (B). An additional example of an electron tomographic view of the curved envelope and its associated structures. (C). A TEM image of the 52 h APF olfactory sensillum after treatment with 2% SDS. Scale Bar: 100nm (A), 50 nm (B, C).

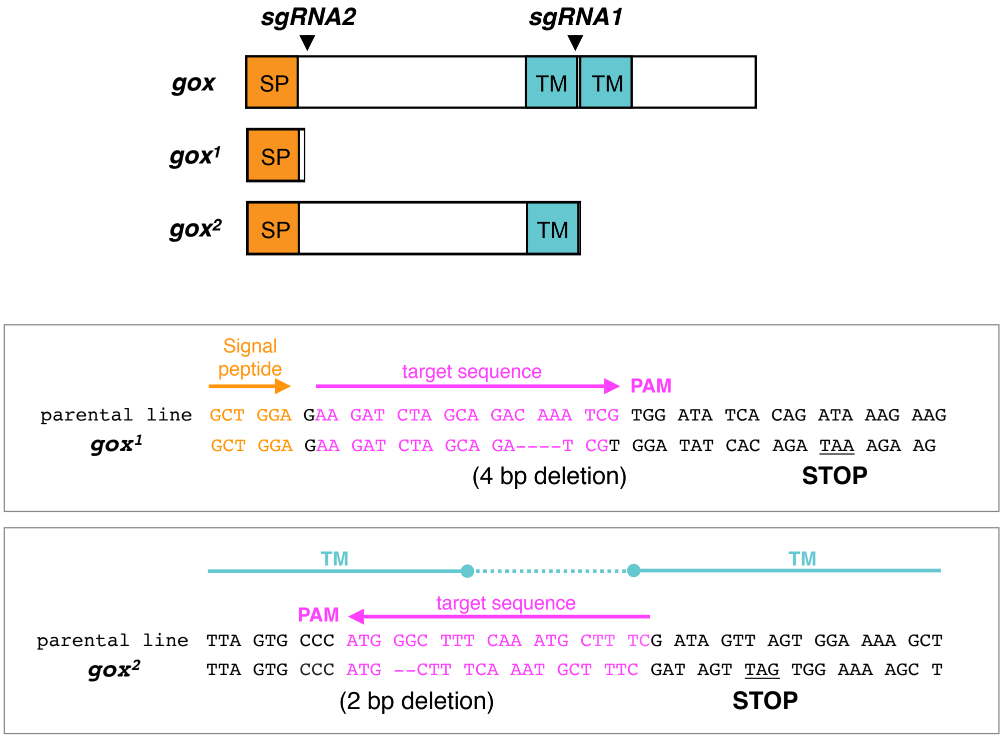


**Figure S2. CRISPR/Cas9-induced *gore-tex/Osi23* mutations. Related to Figure 3**

sgRNA2 targeted to the sequence immediately downstream of the signal-peptide sequence generated *gox*$^2$, and sgRNA2 targeted to the sequence between the two putative transmembrane sequences generated *gox*$^1$.

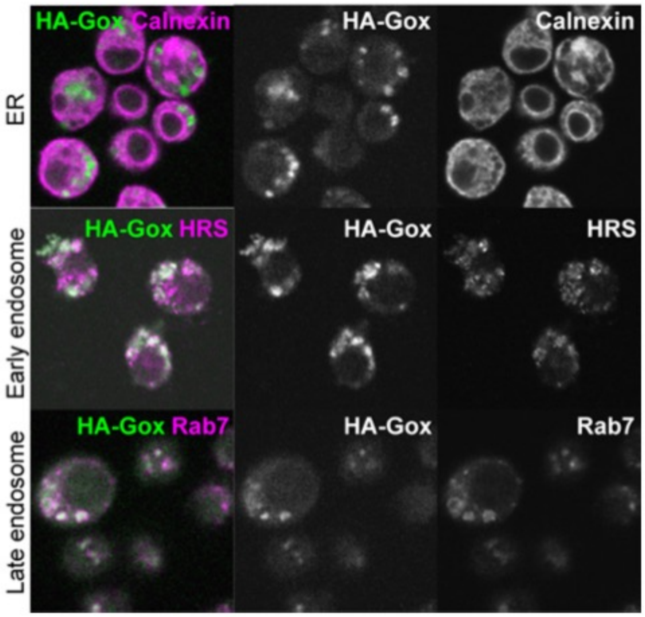


**Figure S3. Analysis of Gox protein in S2 cells. Related to Figure 3**

HA-Gox was colocalized with HRS (early endosomes) and Rab7 (late endosomes) but not with Calnexin (endoplasmic reticulum [ER]). Bar: 10 µm.

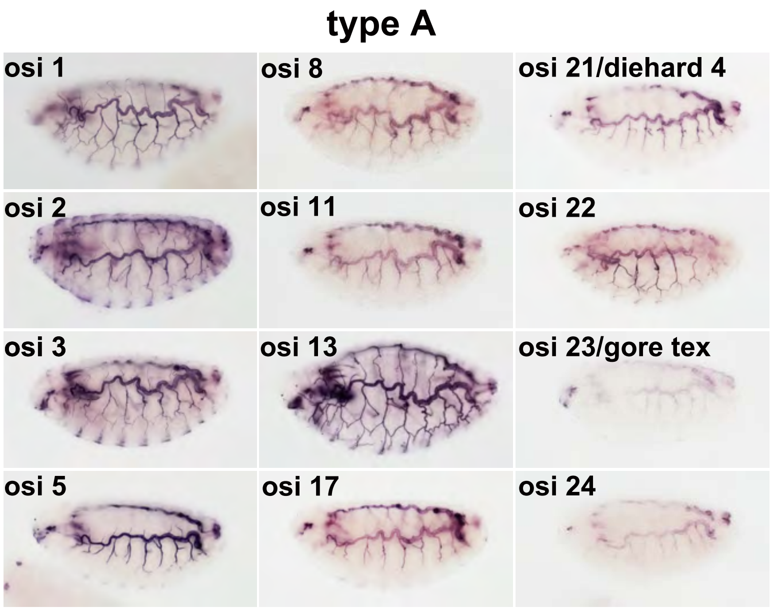
type A
osi 1
osi 8
osi 21/diehard 4
osi 2
osi 11
osi 22
osi 3
osi 13
osi 23/gore tex
osi 5
osi 17
osi 24

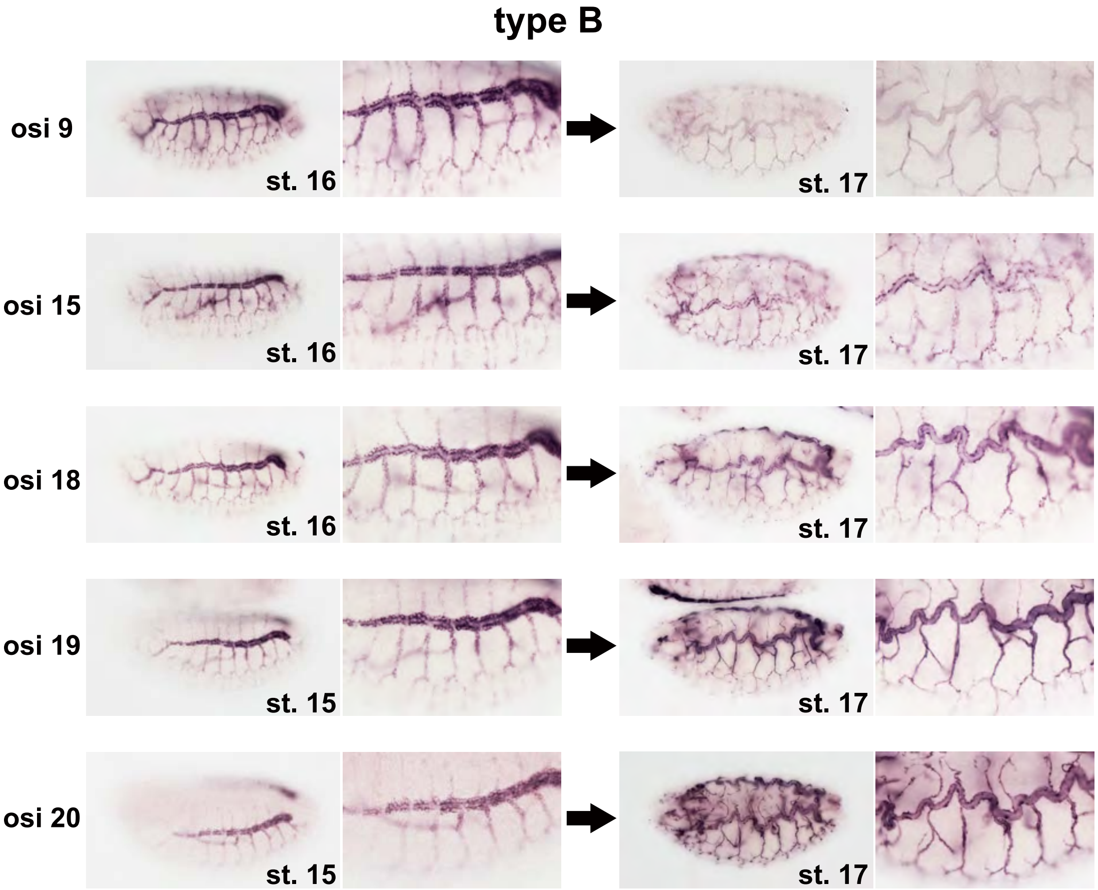
type B
osi 9
st. 16
st. 17
osi 15
st. 16
st. 17
osi 18
st. 16
st. 17
osi 19
st. 15
st. 17
osi 20
st. 15
st. 17

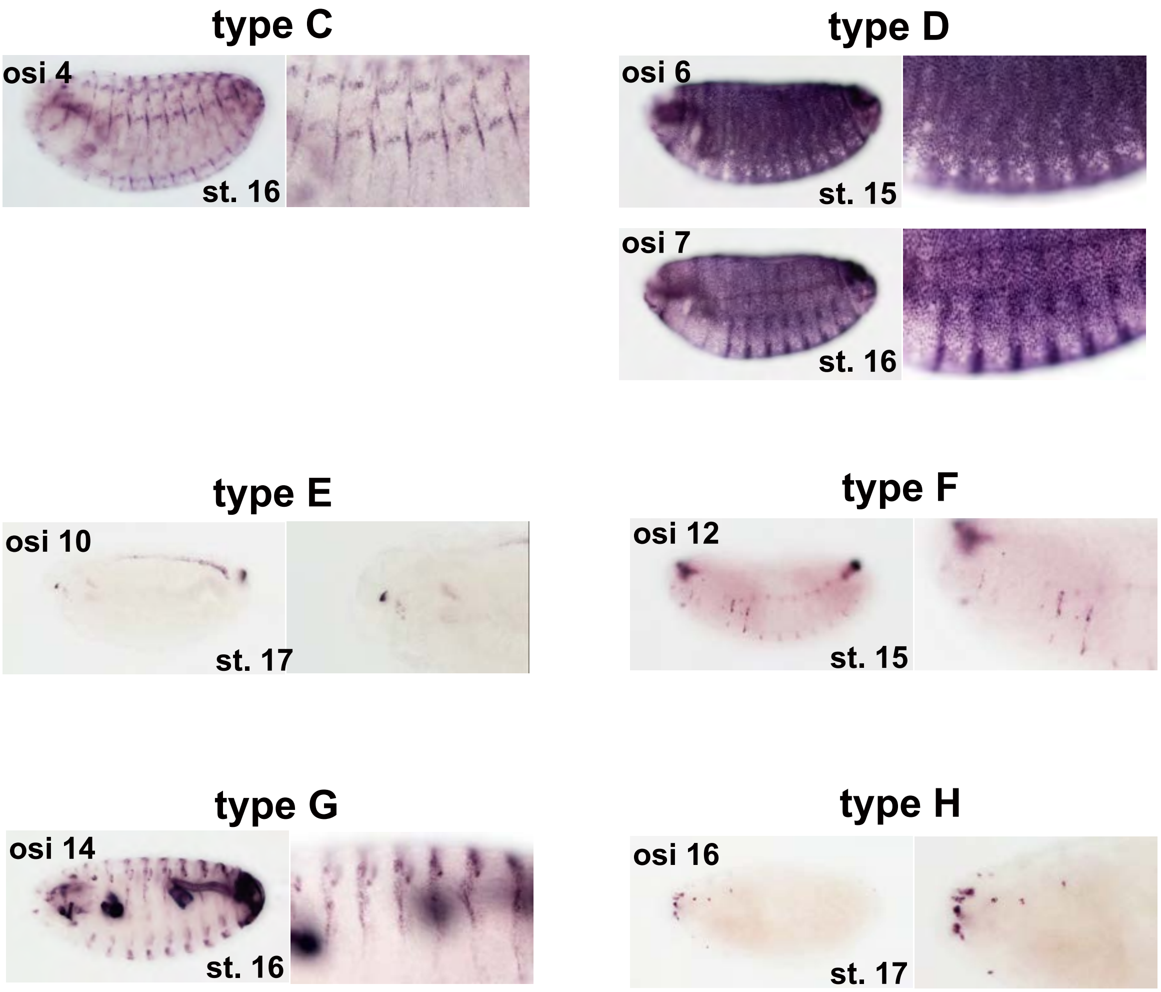


**Figure S4. Eight types of *Osi* gene expression patterns in the embryo. Related to Figure 4.**
Eight types of *Osi* gene expression patterns in the *Drosophila* embryo. Type A genes (*Osi1, Osi2, Osi3, Osi5, Osi8, Osi11, Osi13, Osi17, diehard4/Osi21, Osi22, gore-tex/Osi23, Osi24*) were expressed in the trachea, head skeleton, and posterior spiracle to various degrees. Type B genes (*Osi9, Osi15, Osi18, Osi19, Osi20*) were detected in the cytoplasm of tracheal cells, except for fusion cells at stage 17. At stage 17, the mRNA signal became sharply concentrated in the apical plasma membranes. Type C (*Osi4*): epidermal stripes. Type D (*Osi6, Osi7*): Broad epidermal expression. Type E (*Osi10*): 4-cell cluster in the anterior pharynx. Type F (*Osi12*): anterior pharynx and posterior spiracle. Type G (*Osi14*): foregut, hindgut, and salivary gland duct. Type H (*Osi16*): a small number of epidermal cells.

<u>Additional text on the expression pattern of the 24 *Osi* family genes</u>

According to the modENCODE temporal RNA expression data available from Flybase (http://flybase.org), the expression levels of the *Osi* genes vary from no/extremely low to extremely high in the 14 h-16 h embryo [RPKM (Reads per kilobase of exon model per million mapped reads) value between 0 to 2218, Table S1]. However, our whole-mount in situ hybridization detected tissue-specific mRNA expressions for all 24 genes, including 6 genes classified as “No/Extremely low expression” (*Osi1, Osi8, Osi11, Osi21, Osi22, Osi23*), starting from stage 14 or later. The expression patterns shown in Fig. S4 were highly dynamic and could be categorized into 8 types (A to H, Fig. S4). *Osi* gene expression was most frequently detected in the trachea. The head skeleton, epidermis, fore- and hindgut, and salivary duct, were found to express *Osi* genes. Taken together, most, if not all, of the cuticle-producing epidermal tissues expressed one or more *Osi* gene. Epidermal tissues can thus be considered as a mosaic of *Osi* expression domains.

Type A was the largest category, including 12 genes, and was characterized by its expression in all tracheal cells, and the head skeleton, posterior spiracle, and denticle cells. The mRNAs were mostly associated with the apical plasma membrane and the tip of denticles. Type B (5 genes) was also characterized by its prominent tracheal expression, but its expression was excluded in the two fusion cells in each of the anastomosis site of the dorsal trunk. The mRNAs were mainly localized to all of the cytoplasmic regions of stage 16 trachea, but became sharply concentrated at the apical plasma membrane at stage 17. Type C (*Osi4*) was uniquely expressed in the cells forming segment boundaries. Type D (*Osi6* and *Osi7*) showed strong and broad expression in the epidermis that persisted until stage 16. The mRNA expression was excluded in some parts including the sense organs, posterior spiracle, anterior spiracle, and positions of the leg and wing imaginal discs. The epidermal expression was down-regulated at stage 17, and the expression in the trachea and posterior spiracle became prominent. Foregut and hindgut expressions were also observed. Type E (*Osi10*) was characterized by its very limited expression in 4 cells located in the anterior end of the pharynx. Type F (*Osi12*) showed expression in the anterior pharynx, head sense organs, and posterior spiracle including the filzkölper. In the trachea, the *Osi12* expression was initially high in the dorsal trunk fusion cells, and at stage 17 it became expressed in all tracheal cells. Type G (*Osi14*) showed strong foregut and hindgut expression, and expression in the salivary gland duct and epidermal stripes. Type H (*Osi16*) was expressed in pairs of cells at the position where the leg and wing imaginal discs are connected to the epidermis. Signal clusters in the head were also observed.

| | | expressed tissues | | | | | expressed stage | | | | mRNA level |
|---|---|---|---|---|---|---|---|---|---|---|---|
| gene | type | trachea | head, mouth parts | cuticle | gut | others tissue | st.14 | st.15 | st.16 | st.17 | |
| Osi1 | A | apical surface, ps | AntSO, ci, lr, eps, VA | db | ph, es | | | | | | 1 |
| Osi2 | A | apical surface, ps | AntSO, ci, lr, eps, VA | whole body(week), db | ph, es, hg | | | | | | 5 |
| Osi3 | A | apical surface, ps | AntSO, ci, lr, eps, VA, DBr, DA, MH | db | ph, es | | | | | | 4 |
| Osi4 | C | ps | | striped pattern (segment gr | ph ,es, pv,hg(w | pns | | | | | 2 |
| Osi5 | A | apical surface, ps | AntSO, ci, lr, eps, VA | db | ph, es | signal clusters are shown in head. | | | | | 4 |
| Osi6 | D | apical surface(st.17), ps | DBr, DA, | expressed in whole epidermis expect for head part and pns. db | ph, es, pv, | | | | | | 8 |
| Osi7 | D | apical surface(st.17), ps | AntSO, ci, lr, eps, VA | expressed in whole epidermis expect for head part and pns. | ph, es, pv | | | | | | 8 |
| Osi8 | A | apical surface, ps | AntSO, ci, lr, eps, VA | | es | | | | | | 1 |
| Osi9 | B | cytoplasm(st14-16) and apical (st.17), ps | sd | | | | | | | | 7 |
| Osi10 | E | | | | | signals are shown in only few cells (4 cells?) at lower ph. | | | | | 2 |
| Osi11 | A | apical surface, ps | AntSO, lr, eps, VA | db | ph, es | | | | | | 1 |
| Osi12 | F | ps | AntSO, lr, eps, VA | | | signal clusters in head. | | | | | 6 |
| Osi13 | A | apical surface, ps | AntSO, lr, eps, VA, dbr, MH | db | ph, es | | | | | | 2 |
| Osi14 | G | apical surface(st17), ps | sd | striped pattern | hg, pv | pns, base of anal pad | | | | | 7 |
| Osi15 | B | cytoplasm(st14-16) and apical (st.17), ps | AntSO,lr(week), eps (week) | | | | | | | | 7 |
| Osi16 | H | | | | | clusters of cells in head (lbd, clb.?). two cell signals in T1-T3 correspond to the positions where imaginal discs (wd,hd,ld) are connected to the epidermis. some signal around ps. | | | | | 4 |
| Osi17 | A | apical surface, ps | AntSO, lr, eps, VA | db | ph | | | | | | 5 |
| Osi18 | B | cytoplasm(st14-16) and apical (st.17), ps | | | | | | | | | 7 |
| Osi19 | B | cytoplasm(st14-16) and apical (st.17), ps | AntSO, lr, eps, VA, DBr, DA | db | | | | | | | 7 |
| Osi20 | B | cytoplasm(st14-16) and apical (st.17), ps | AntSO, lr, eps, VA, DBr, DA | db | ph | | | | | | 7 |
| diehard4/Osi21 | A | apical surface, ps | AntSO, lr, eps, VA | db | ph, es | | | | | | 1 |
| Osi22 | A | apical surface, ps | AntSO, lr, eps, VA | db | ph, es | | | | | | 1 |
| gore-tex/Osi23 | A | apical surface, ps | AntSO, lr, eps | | | | | | | | 1 |
| Osi24 | A | apical surface, ps | AntSO, lr, eps, VA, DBr, DA | db | pv | | | | | | 5 |

**Table S1. Summary of the 24 *Osi* gene expression. Related to Figure 4.**

Abbreviation: ps: posterior spiracle, AntSo: antennal sense organ, lr: lareum, eps: epistomal sclerite, VA: ventral arm, DBr: dorsal bridge, DA: dorsal arm, MH: mouth hook, ci: cirri, sd: salivary grand duct cell, db: denticle belt, ph: pharynx, es: esophagus, pv: proventriculus, hg: hindgut, ldb: labial disc, clb: clypeolabral disc, wd: wing disc, hd: haltere disc, ld: leg disc, ko: Keilin's organ.

mRNA expression levels at 14-16 h after egg laying (stage 16) from the temporary expression data of modENCODE is shown in the rightmost column. Eight categories based on the RPKM (Reads per kilobase of exon model per million mapped reads of RNA-seq data) values are shown.

Bin 1: No/Extremely low expression (0)
Bin 2: Very low expression (1-9), percentiles 1-25, approximately.
Bin 3: Low expression (10-150), percentiles 26-50, approximately.
Bin 4: Moderate expression (151-2500), percentiles 51-75, approximately.
Bin 5: Moderately high expression (2501-7500), percentiles 76-85, approximately.
Bin 6: High expression (7501-25000), percentiles 86-95, approximately.
Bin 7: Very high expression (25001-100000), percentiles 96-99, approximately.
